# Looking backward for the future: long-term population recovery, habitat associations, and future climatic vulnerability of the critically endangered Sino-Mongolian beaver (*Castor fiber birulai*) in China

**DOI:** 10.64898/2026.02.07.704560

**Authors:** Wenwen Chu, Yuanbao Du, Roberto Salguero-Gómez, Yingjie Qi, Chi Ma, Wenxu Lan, Xiaoyun Li, Rukeya Abulimit, Fangling Zheng, Ziyu Liu, Yuting Gao, Huanxin Liu, Chengbin He, Kai Li, Hongjun Chu

**Author notes:** These authors equally contributed to this manuscript (Wenwen Chu, Yuanbao Du). Corresponding authors: Yuanbao Du; Kai Li; Hongjun Chu.

## Abstract

Despite the successful population recovery of the Eurasian beaver (*Castor fiber*) across much of Eurasia, its subspecies, the Sino-Mongolian beaver (*C. f. birulai*), remains critically endangered, with an estimated population of approximately 1,500 individuals confined to a small number of isolated and fragmented refugia along the China–Mongolia border. Effective conservation of this highly threatened subspecies requires a holistic perspective that integrates constraints on population dynamics, habitat associations, and future climatic vulnerability. Here, we combined systematic annual field surveys conducted between 2003 and 2023 with historical survey records from 1975 to 1989 in northern Xinjiang, China, to synthesize long-term spatiotemporal population dynamics, evaluate habitat preferences based on nine local environmental variables, and assess future climatic vulnerability using ensemble species distribution models (SDMs) under projected climate change scenarios. We detected a significant and phased population recovery, with beaver colony numbers increasing from 27 (approximately 100 individuals) in 1975 to 227 (681–908 individuals) in 2023. This recovery closely corresponded with major conservation milestones, including the establishment and upgrading of nature reserves, strengthened legislative protection, and enhanced multi-stakeholder collaboration. Habitat analyses further indicated that the Sino-Mongolian beaver preferentially occupied areas characterized by minimal anthropogenic disturbance and stable hydro-geomorphic conditions. Critically, SDM projections revealed that only 14% of the current study area presently exhibits high climatic suitability, and these highly suitable habitats are expected to disappear entirely by the 2050s. Together, our findings provide a comprehensive overview of the historical population recovery and conservation trajectory of the Sino-Mongolian beaver in China, and offer robust scientific support for developing adaptive management strategies in the face of ongoing climate change and increasing human pressures.

## Introduction

As one of only two extant species within the *Castor* genus, the Eurasian beaver (*Castor fiber*) is widely recognized as a keystone species and an exceptional ecosystem engineer in freshwater systems across Eurasia continent [1–3]. Through dam construction, channel excavation, and wetland creation, Eurasian beavers play a vital role in maintaining ecological stability and enhancing biodiversity [1–3]. However, centuries of intensive overhunting for fur, meat, and castoreum have drastically reduced their numbers to approximately 1,200 individuals, confined to a few isolated refugia [2–9]. Since the early 20th century, extensive conservation efforts including the protect areas establishment and reintroduction programs have jointly resulted in a remarkable population recovery, particularly in Europe [2, 4, 7, 8, 10, 11]. By 2020, the Eurasian beaver population had rebounded to more than 1.5 million individuals [2, 7], which also led to a *Least Concern* status classified by the International Union for Conservation of Nature (IUCN) [12]. Nonetheless, human-deduced challenges such as habitat loss and fragmentation, competition and disturbance by domestic animals continue to threaten certain subspecies, especially the Sino-Mongolian beaver (*C. f. birulai*) [12].

As one of recognized subspecies of the Eurasian beaver [13], the Sino-Mongolian beaver remains *Critically Endangered*, with a total of roughly 1,500 individuals restricted to limited areas in China and Mongolia [1, 7, 9, 14]. Historically, the beaver population in China ever declined to fewer than 100 individuals in the late 1970s, which were confined to the mainstem and tributaries of the Ulungur River in northern Xinjiang [1, 15]. In the early 1980s, China government established a province-level nature reserve (Xinjiang Burgen Beaver Nature Reserve) to protect the Sino-Mongolian beaver and over 5,000 ha of core beaver habitat in Qinghe County. Subsequently, in 1989, the promulgation of the Wildlife Protection Law of China and the release of the National Key Protected Wildlife Checklist designated the Sino-Mongolian beaver as a national first-class protected species. These institutional efforts had initially facilitated a substantial population recovery with colony counts rising to 165 in the late 1980s [16]. Subsequently, the upgrade of the nature reserve to a national level (Xinjiang Burgen Beaver National Nature Reserve) in 2013 underscored an intensified conservation commitment, which contributed to the relative stability in beaver population with about 130-170 colonies before 2018. More recently, a series of collaborative conservation programs were launched in 2019, through integrating efforts of government, social donations, conservation societies, and local communities under the Shining Stone Community Actions (SSCA). Nevertheless, there have been more than 40 years since the original conservation actions in early 1980s. How these institutional and collaborative conservation efforts acted on the beaver population recovery has not been systematically reviewed by now with detailed population data. Besides, as the population recovers within this human-settled landscape in this arid area, how beaver adapt to local human disturbance environment has not been systematically reviewed.

In addition to human disturbance, climate changes are also recognized to pose escalating and significant threats to beaver population persistence through their effects on thermoregulation, forage availability, and life-history processes [17, 18]. Generally, climate changes affect beaver population by altering temperature- and rainfall-driven plant phenology and forage quality and availability, which influence annual body weight and reproductive success [18]. Increasing climatic variability, particularly higher temperature variance and precipitation variability—further influences vital rates, with greater variability generally reducing survival (especially of kits, juveniles, and dominant adults) and shaping recruitment patterns. Overall, both shifts in climate means and increases in variability play key roles in regulating beaver population dynamics through their effects on thermoregulation, forage availability, and life-history processes. As shifts in hydrothermal regimes have intensified the occurrence of extreme climate events including droughts and floods, which contribute to higher mortality for beavers through pronounced fluctuations in river hydrology [17]. Climate change affects Eurasian beaver populations by altering temperature- and rainfall-driven plant phenology and forage quality, ultimately reducing body weight and reproductive success. Although previous studies have ever examined population status [1, 16], habitat preference [19, 20], and behavioral ecology [21, 22], there remains a pressing need for a comprehensive synthesis integrating long-term population dynamics, habitat selection, and climatic vulnerability under ongoing climate change.

To address this gap, by integrating confidential data extracted from previous studies (1975-1989) and our systematic annual field surveys (2003-2023) across the main distribution areas in China, this study aims to: (1) reconstruct the long-term spatiotemporal population dynamics of the Sino-Mongolian beaver and assess the effectiveness of previous conservation efforts; (2) evaluate its habitat association in relation to nine local environmental variables; and (3) employ ensemble species distribution modeling to assess climatic vulnerability under climate change scenarios. These findings are promising in that they provide a comprehensive picture of the history of Sino-Mongolian beaver population recovery and conservation efforts in China, and offer scientific support for developing future management strategies.

## Methods and Materials

### Study area

The study area encompasses the main distribution area of the Sino-Mongolian beaver in China, covering the mainstream and tributaries of the Ulungur River in northern Xinjiang (87.16–91.07°E, 45.81–47.39°N; Fig. 1). This region, adjacent to the Eastern Altai Mountains and the China-Mongolia border, features terrain spanning 465 m to 3,530 m in elevation. The climate is typical continental cold-temperate, characterized by winters lasting approximately six months and a frost-free period of 67–115 days.Recorded temperatures range from −49.7°C in winter to 34.3°C in summer. In this arid region, annual precipitation is equivalent to only 10% of the annual evaporation, with approximately 33% of the annual precipitation falling as snow. During winter, snow cover typically reaches a depth of 80 cm, and the frozen soil layer extends to 70 cm [1].

**Fig. 1.**
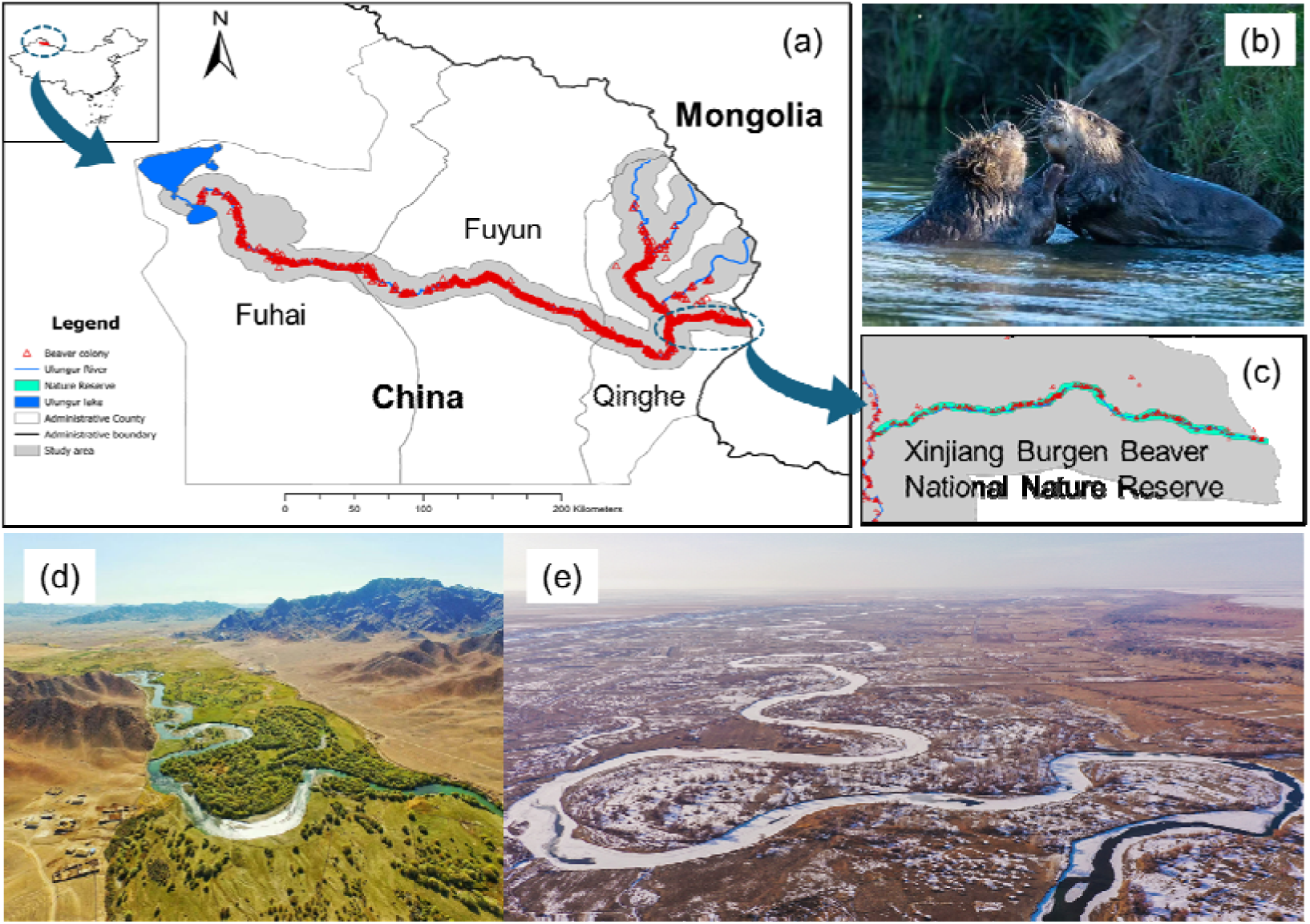
Overview of study area and the Sino-Mongolian beaver. (a) Map showing the location of the study area in northern Xinjiang, China. (b) Two Sino-Mongolian beavers. (c) Spatial extent of the Xinjiang Burgen Beaver National Nature Reserve. (d) and (e) show representative landscapes of the beaver habitat in summer and winter, respectively.

### Field survey

We conducted 14 comprehensive annual field surveys for the Sino-Mongolian beaver over a 21-year period (2003–2023). Field survey was carried out between late October and December within a 5-km buffer zone along the mainstream and three tributaries of the Ulungur River, encompassing all associated water bodies (e.g., rivers, streams, ponds, wetlands, and agricultural channels; Fig. 1), which covered approximately 14,967 km². Field survey teams, each comprising at least three investigators, systematically identified beaver nests, dams, and food piles (Fig. S1); for this study, an active beaver nest was considered indicative of an active beaver colony. We recorded the geographic coordinates (latitude, longitude, and elevation) of each active nest using a handheld GPS device (UniStrong UG908) and referenced the locations using Ovital Map (https://www.ovital.com/).

### Population dynamics across space and time

The population size of beavers is typically estimated by integrating colony number and the size of food cache for each beaver colony [1, 23]. However, during winter periods, snow and ice can partially cover food caches, leading to unavoidable under-or overestimation of volume that varies significantly across investigators. To mitigate this systematic variability, we characterized the population dynamic in this work using colony count and colony density, rather than estimating individual numbers directly. Previous studies reported that each beaver colony contains 3 to 4 individuals with an average of 3.8 [1, 16], and these values were adopted as the lower, upper, and average parameters when estimating the total beaver population.

#### Population dynamic model: colony count (2003-2023)

Given the more than 20-year gap between our data and those reported previously (Fig. 2), our population dynamic models were fitted exclusively using colony counts obtained from our annual field surveys conducted from 2003 to 2023. To better characterize demographic dynamics across space and time, we divided the entire study area into two parts based on digital and aquatic conditions: the upstream area including the mainstream of the Ulungur River and three tributaries in Qinghe county, and the downstream area including the mainstream of the Ulungur River in Fuyun and Fuhai counties [1] (Fig. 1a). Additionally, in the upstream area, we also characterized the beaver colonies within the nature reserve alone to evaluate its conservation efficiency.

**Fig. 2.**
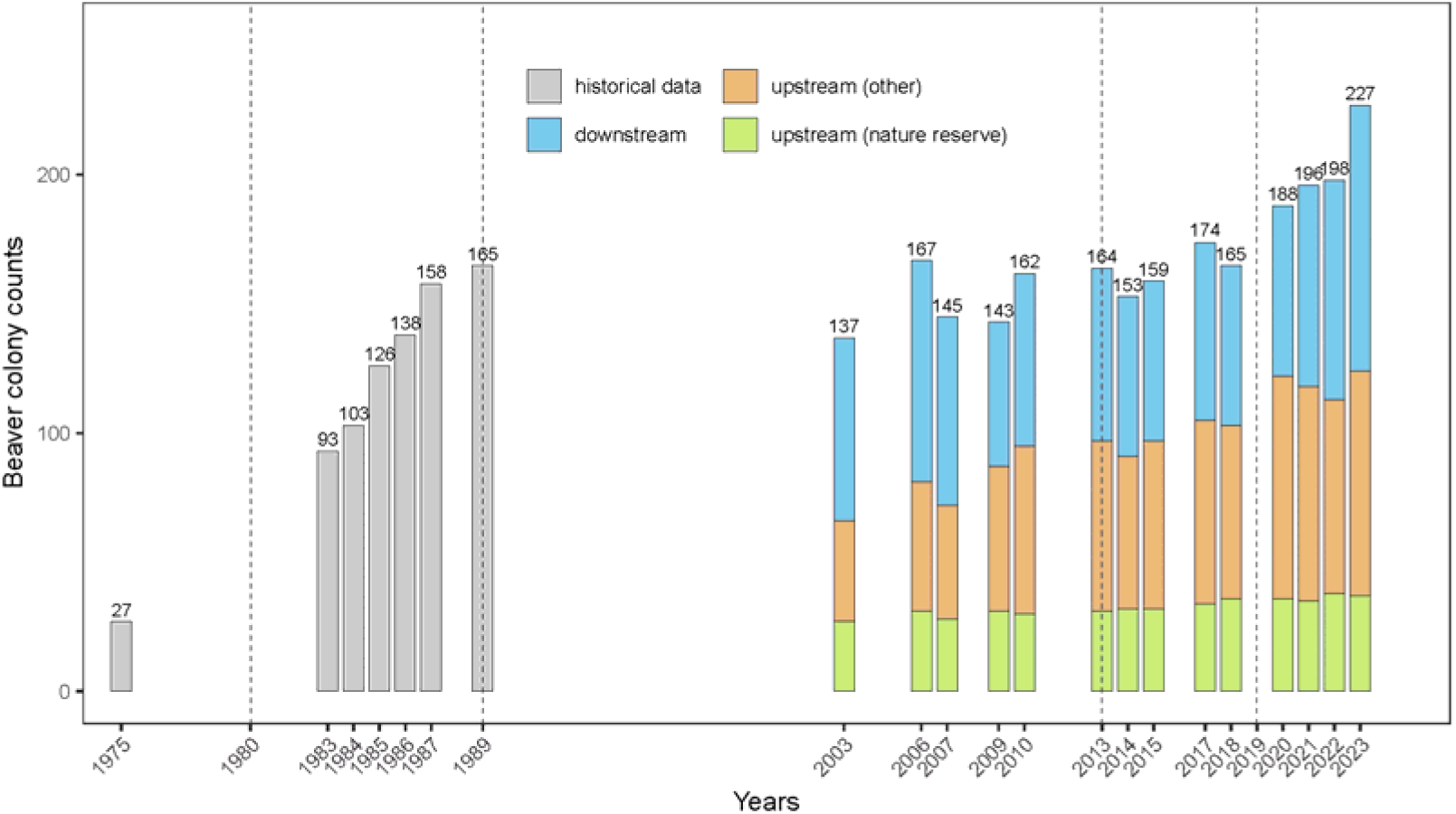
Historical population dynamics (colony counts) of the Sino-Mongolian beaver in northern Xinjiang, China (1975–2023). Historical beaver colony data (1975-1989) are shown by light gray bars (adapted from Shao, 1990). Annual beaver colony counts collected in this study are represented by colored cumulative bars: orange denotes upstream sites (excluding the nature reserve), grass green denotes sites within the nature reserve, and steel blue denotes downstream sites. Key conservation milestones are marked by dashed lines (from left to right): (1) establishment of the nature reserve at the provincial level in 1980; (2) promulgation of Wildlife Protection Law of China and the beaver’s listing as a *First-Class* Key Protected Wildlife of China in 1989; (3) upgrading the reserve to a national status in 2013; and (4) conducting collaborative conservation projects by integrating government, conservation society, and local communities in 2019.

To model the annual beaver colony count over the time of investigation since 2003, we applied three approaches: the exponential growth model, the polynomial regression model, and the logistic growth model. To enhance model convergence, the colony count (response variable) and time (independent variable) were both log-transformed.

We fitted the exponential growth model using non-linear least squares (NLS) for the whole study area, upstream, downstream, and the nature reserve separately, based on the equation:

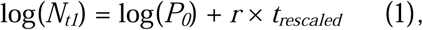

where *N_t1_*is the colony number at year *t*, *t_rescaled_* is the rescaled year *t*, *P_0_* is the initial colony number at first year (2003), and *r* is the intrinsic growth rate, which was assumed to be 0.05.

Next, we fitted polynomial regression model using the equation:

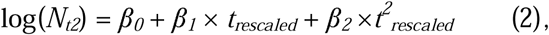

where *N_t2_*is the colony number at year *t*, *t_rescaled_* is the rescaled year *t,* β*_0_* is the intercept, β*_1_*and β*_2_* are the coefficients of the linear and quadratic components, respectively.

Finally, the logistic growth model was also fitted using the NLS method with the “port” algorithm, based on the equation:

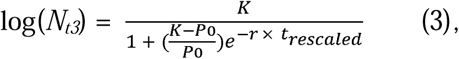

where *N_t3_* is the colony number at year *t, K* represents the carrying capacity (set as maximum colony *1.2), *P_0_* is the initial colony number at first year (2003), and *r* is the intrinsic growth rate (0.05). The performance of all three demographic models was ultimately assessed using the Akaike Information Criterion (AIC).

#### Spatiotemporal dynamic of colony density (2003-2023)

We applied the *Kernel Density* function to describe the spatial pattern of beaver colony density across the study area. To further characterize temporal variation in density, we divided our 14 annual field surveys into five stages, with each stage containing three years of beaver nest distribution data. Specifically, the stages were defined as: stage 1 (2003, 2006, and 2007), stage 2 (2009, 2010, and 2013), stage 3 (2014, 2015, and 2017), stage 4 (2018, 2020, and 2021), and stage 5 (2021, 2022, and 2023). Given the limited number of survey years, the data for 2021 was necessarily shared between stage 4 and stage 5 to maintain consistency in calculating colony density and temporal variation. Following these temporal divisions, we calculated the beaver colony density for each stage and quantified their temporal variation across adjacent stages. The colony densities and their temporal variations were then mapped using optional parameters within ArcGIS Pro version 3.0.1 (https://www.esri.com/).

### Habitat associations

We used nine habitat variables relating to anthropogenic disturbance, hydrological conditions, and biotic/abiotic factors to determine the dominant drivers of Sino-Mongolian beaver habitat association. Anthropogenic disturbance variables included three measures: the distance of the beaver nest to the (1) nearest road, (2) nearest human settlement, and (3) nearest artificial river dam. The hydrological variable was (4) the water depth measured 2 meters from the riverbank near beaver nest. The biotic and abiotic variables were (5) elevation of the nest, (6) forest canopy within a 50-meter radius of the nest, (7) the nest height above the riverbank, (8) the horizontal distance to the riverbank, and (9) the Normalized Difference Vegetation Index (NDVI) at the nest location. Elevation, nest height, and canopy were recorded during field surveys. For the distance metrics, we obtained spatial vector data for roads, dams, and human settlements. We utilized global datasets for this purpose: the Global Human Settlement Layer (GHSL, accessed on October 20th, 2024), the Global Lakes and Wetlands Database (GLWD, accessed on October 20th, 2024), and the GRIP global roads database [24] (accessed on October 20th, 2024). Artificial dam locations were identified and recorded during field surveys. We then calculated the nearest distance metrics using the *Proximity Analysis* function in ArcGIS and extracted these distance-based variables according to the geographic coordinates of each beaver nest. The NDVI data was extracted from the 1-kilometer resolution MOD13Q1 remote layer, downloaded from the Geographic Remote Sensing Ecological Network Platform (http://www.gisrs.cn/index.html, accessed in October 2024). Finally, to characterize habitat association, we performed a Principal Component Analysis (PCA) on the nine variables and quantified their relative importance within each PCA component.

### Future risk of habitat loss

To quantify current and future habitat suitability, we performed species distribution modeling (SDM) using three well-established methods known for their robust performance and ability to handle non-linear relationships: Maxent, Generalized Linear Models (GLM), and Random Forest [25]. We generated an ensemble prediction of beaver distribution by equally weighting the results of these individual models [26, 27].

To ensure SDM robustness, we compiled species occurrence data from two sources: over 2,300 records of Sino-Mongolian beaver colony locations collected from 14 annual field surveys from 2003 to 2023, and global records of the Eurasian beaver collected from the GBIF database (https://www.gbif.org/, accessed on January 3, 2025) using the *occ_download* function in *rgbif* package [28] with “*Castor*” and “*fiber*” as keywords in data retrieval (https://doi.org/10.15468/dl.g6h7u2). We only retained the records with detailed geographical information (longitude and latitude) and further conducted data cleaning using the *clean_coordinates* function in *CoordinateCleaner* package [29]. For occurrence data outside our study area, we conducted data thinning using *spThin* package [30] to retain only one randomly selected point per 5-kilometer square.

We included climatic and anthropogenetic disturbance variables in the SDMs. To minimize collinearity (r ≤ 0.7) among climatic variables, we selected six Bioclimatic variables recognized as important factors influencing beaver life history [17]: mean temperature diurnal range (Bio2), temperature isothermality (Bio3), temperature seasonality (Bio4), min temperature of coldest month (Bio6), annual precipitation (Bio12), and precipitation of coldest quarter (Bio19). According to the results of habitat association, we included three anthropogenic disturbance variables: the nearest distance to the river, the nearest distance to human settlement, and the nearest distance to the road. For future projections, these three local anthropogenic disturbance variables were assumed to remain consistent with the current situation.

Model performance was validated using Area Under the Curve (AUC) [31] and True Skill Statistic (TSS) [32]. To reduce overfitting, we used k-fold cross-validation (k = 5) and utilized an independent testing dataset. Future habitat suitability predictions were generated under three Shared Socio-economic Pathways (SSPs: SSP126, SSP370, and SSP585) and four Global Climate Models (GCMs: IPSL-CM6A-LR, MPI-ESM1-2-HR, MRI-ESM2-0, and UKESM1-0-LL) for the 2050s (2041-2060), 2070s (2061-2080), and 2100s (2081-2100). Bioclimatic variables for both current (1970-2000) and future periods were downloaded at a 1-kilometer resolution from WorldClim version 2.1 [33] (https://www.worldclim.org/, accessed on October 20, 2024) at 1-kilometer resolution. For future projections, our main results were based on the SSP370 scenario, with SSP126 and SSP585 used for sensitivity analyses.

Population dynamic modeling, PCAs, and SDMs were all conducted under the R language environment [34] using the *usdm* [35], *sdm* [36], and basic packages. Study area, colony locations, colony density, ensemble SDM predictions, and spatiotemporal variations were visualized using ArcGIS Pro version 3.0.1 (https://www.esri.com/).

## Results

### Historical population dynamics

A total of 2,378 active beaver colonies were recorded during our 14 annual field surveys conducted from 2003 to 2023 in north Xinjiang, China. The colony count for the Sino-Mongolian beaver demonstrated a general, though fluctuating, increasing pattern, rising from 137 colonies in 2003 to 227 colonies in 2023 (Fig. 2). Before 2018 (165 colonies), the number of colonies remained relatively stable, exhibiting only a slight general increase. A notably faster rate of increase was observed in the last five years following the initiation of collaborative conservation programs in 2019, with the colony number reaching 188 colonies in 2020 and subsequently 227 colonies in 2023. Based on the average estimate of 3.8 (Min/Max: 3/4) individuals per colony, the estimated population size increased from approximately 520 (Min/Max: 411/548) individuals in 2003 to 862 individuals (Min/Max: 681/908) in 2023.

Across the entire study area, all three demography dynamic models significantly fitted the log-transformed colony numbers and scaled time (p < 0.05), with the polynomial model exhibiting the best fit (AIC = -32.6; R^2^ = 0.794), indicating a general increasing trend (Fig. 3a). When analyzed spatially, colony trends differed between upstream (Qinghe County) and downstream (Fuyun and Fuhai Counties) areas (Fig. 3b-c): the upstream area showed a steady, near-linear increase over time, whereas the trend in the downstream area was distinctly polynomial, characterized by an apparent decrease before 2010 followed by a sharp increase after 2018. The number of colonies within the nature reserve exhibited a fluctuating but generally increasing pattern, which was consistent to that of the upstream area (Fig. 3d).

**Fig. 3.**
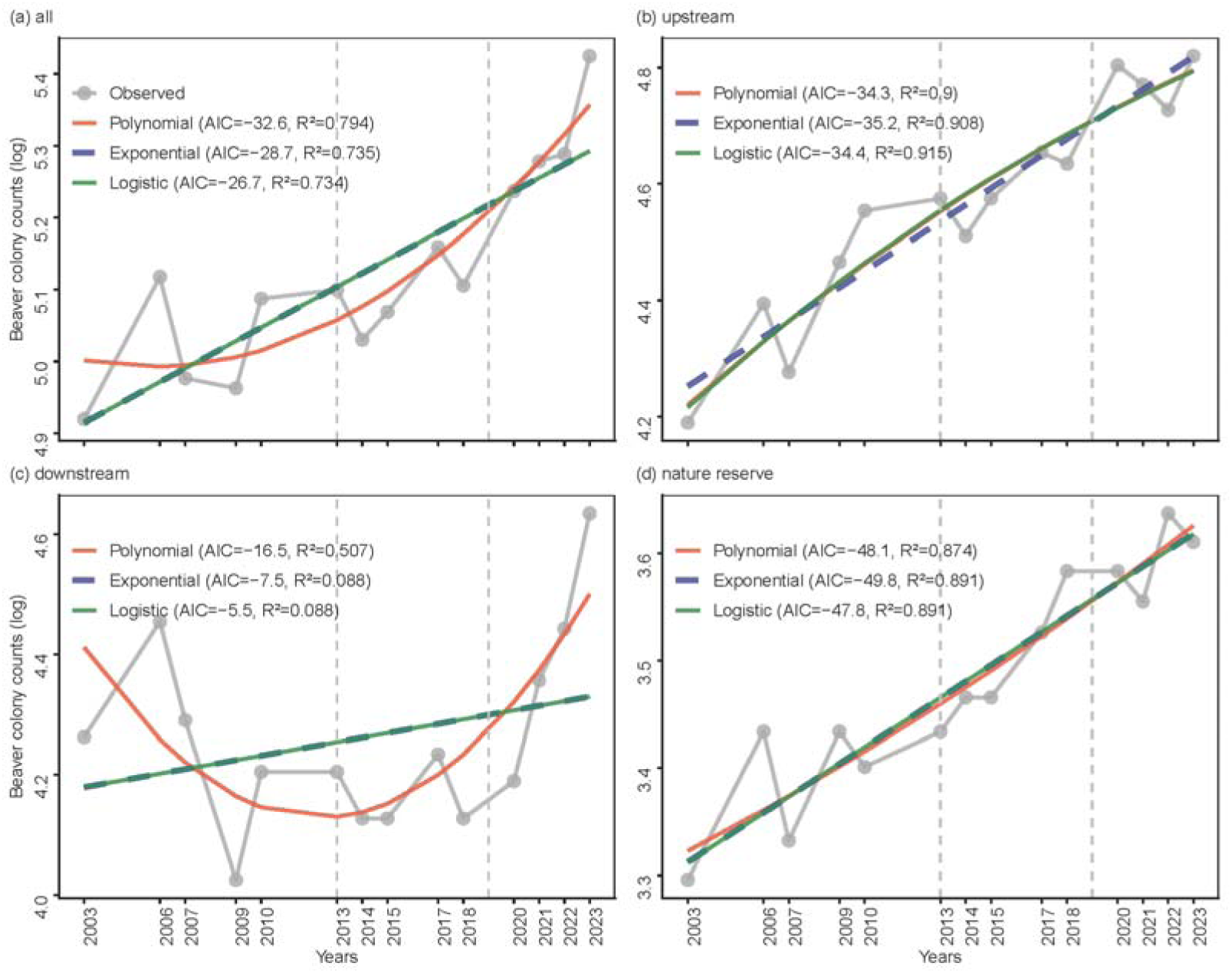
Population dynamic model fitting for beaver colony numbers (2003-2023). Population dynamics were modeled using polynomial (orange), exponential (purple blue), and logistic (green) functions for different spatial categories: (a) the whole study area, (b) upstream areas, (c) downstream areas, and (d) the nature reserve. The Akaike Information Criterion (AIC) values and R^2^ are presented. Dashed vertical lines indicate key conservation milestones: (1) upgrading the reserve to national status in 2013; and (2) conducting collaborative conservation projects by integrating government, conservation society, and local communities in 2019.

The density of beaver colonies generally varied across both space and time (Fig. S2 and 4). Specifically, colonies in the upstream area, including the nature reserve, consistently exhibited a relatively higher density that remained stable across all five temporal stages (Fig. S2 and 4). In contrast, the downstream area had a relatively lower colony density in the early stages (Fig. S2a-b and Fig. 4a-b) but showed a notable increase in the late stages (Fig. S2c-e and Fig. 4c-d).

**Fig. 4.**
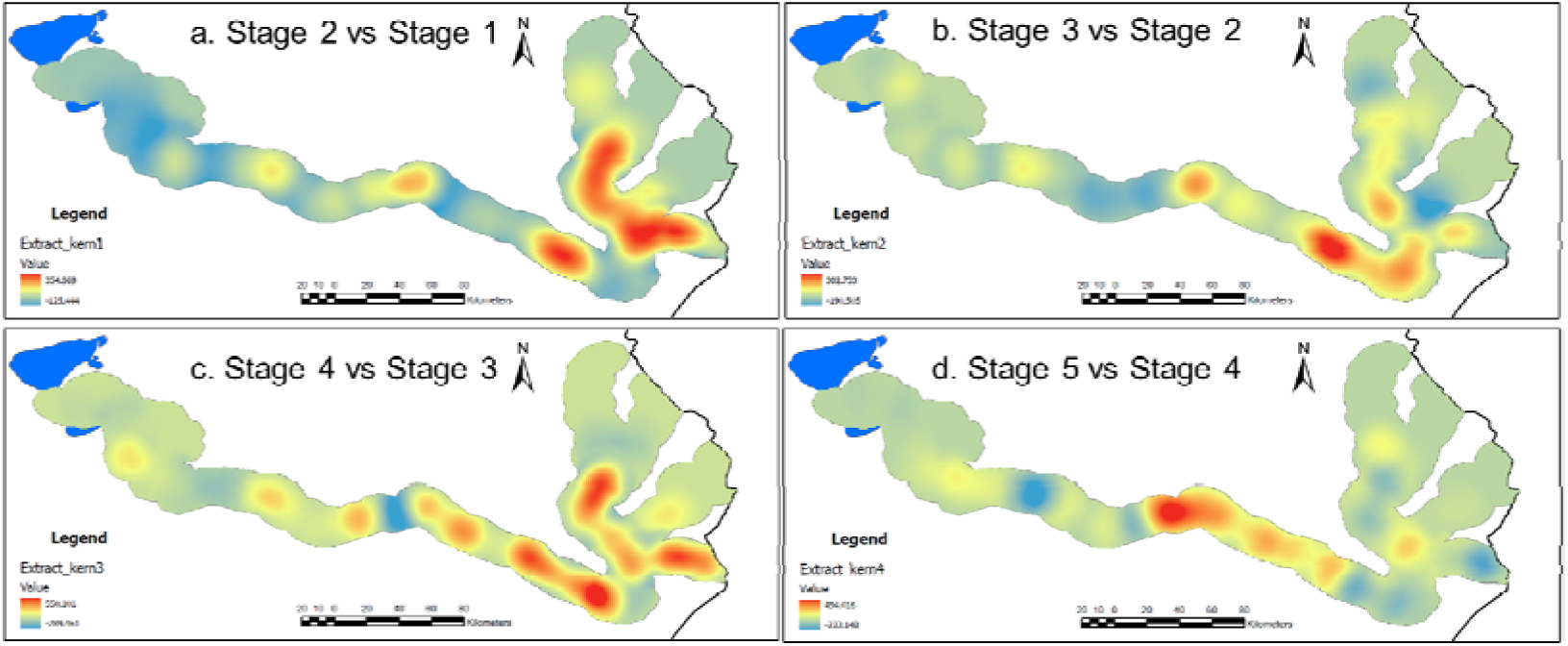
Spatio-temporal variation of beaver colony density across five sequential stages. Each panel illustrates the two-stage temporal dynamics of nest kernel density distribution. Red shading indicates areas where nest density increased during the transition, while light blue shading represents areas where density decreased. Note that the beaver nest distribution data for 2021 was used in the density calculations for both Stage 4 and Stage 5.

### Habitat associations

The results of the PCA suggested that the first four components collectively accounted for 60.2% of the total variance across all nine habitat variables, with PC1 capturing 23.7%, PC2 13.2%, PC3 11.9%, and PC4 11.4% (Fig. 5a-b and Table S1). PC1 was primarily characterized as a gradient of topology and human disturbance, which received the highest contributions from elevation and NDVI, followed by human disturbance variables like distance to artificial dams and human settlements (Fig. 5a, c). Specifically, PC1 exhibited strong positive relationship with distance to artificial dams and human settlements and showed negative relationship with elevation and NDVI (Fig. 5a). By comparison, PC2 displayed as a hydro-geomorphic gradient, demonstrating a positive relationship with the distance to rivers and roads, but a negative pattern with the water depth and nest height (Fig. 5a). The remaining variance was captured by PC3 and PC4, with PC3 being most positively influenced by canopy cover and PC4 being most negatively related to the distance to roads (Fig. 5b-c).

**Fig. 5.**
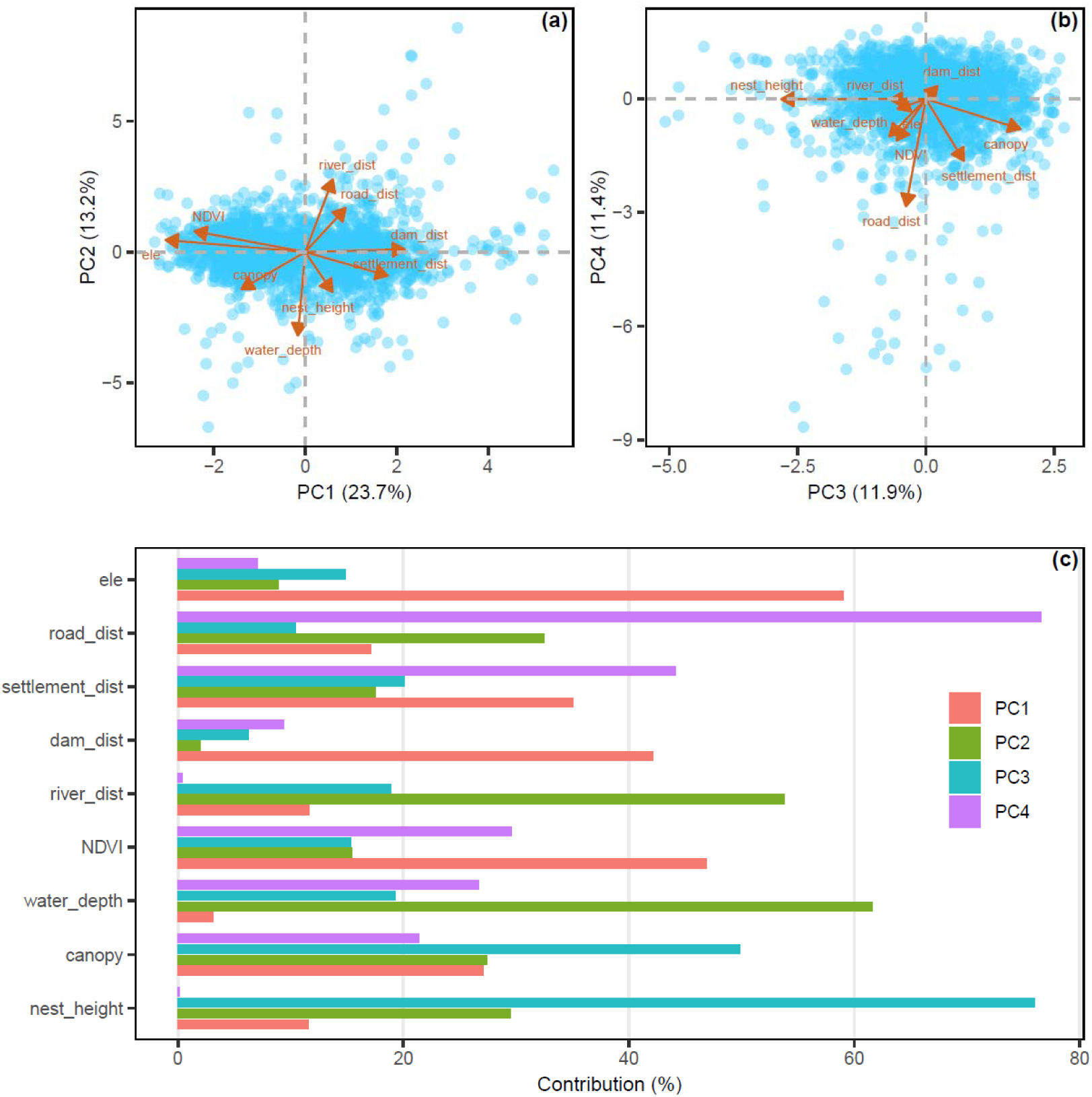
Principal component analysis (PCA) of nine habitat variables. PCA was performed on nine habitat variables within the study area. (a) and (b) illustrate the performance of the nine variables across the PCA components. (c) displays the contribution of each variable to the formation of the respective PCA components. Abbreviations: ele: Elevation; road_dist: Distance to nearest road; settlement_dist: Distance to nearest human settlement; dam_dist: Distance to nearest artificial dam; canopy: Forest canopy cover near the beaver nest; NDVI: Normalized Difference Vegetation Index near the beaver nest; water_depth: Water depth near the beaver nest; nest_height: Height of the beaver nest to the riverbank.

### Risk of climatic suitability decline

The species distribution models (SDMs), which incorporated six climatic and three anthropogenetic disturbance variables, demonstrated stable and robust performance with high values for the AUC (Mean ± SD = 0.997 ± 0.004) and the TSS (Mean ± SD = 0.978 ± 0.019) (Table S2). Based on correlation and AUC tests, five climatic variables (Bio12, Bio3, Bio2, Bio4, and Bio19) were found to have a relatively high contribution to the SDM predictions (Table S3).

The SDM results suggested that there was only 14% of the area now exhibited high climatic suitability (0.8-1.0), while 45% was classified as unsuitable (0-0.2) (Fig. 6a). Under the SSP370 scenario, climatic habitat quality declined significantly (Fig. 6b), resulting in the disappearance of all highly suitable areas in the 2050s, 2070s, and 2100s. At the same time, the proportion of unsuitable areas (0-0.2) rose to 56%, 79% and 82% in the 2050s, 2070s, and 2100s, respectively (Fig. 6b-d). The alternative scenarios SSP126 and SSP585 showed similar trends to those under SSP370 scenario (Fig. S4-S5).

**Fig. 6.**
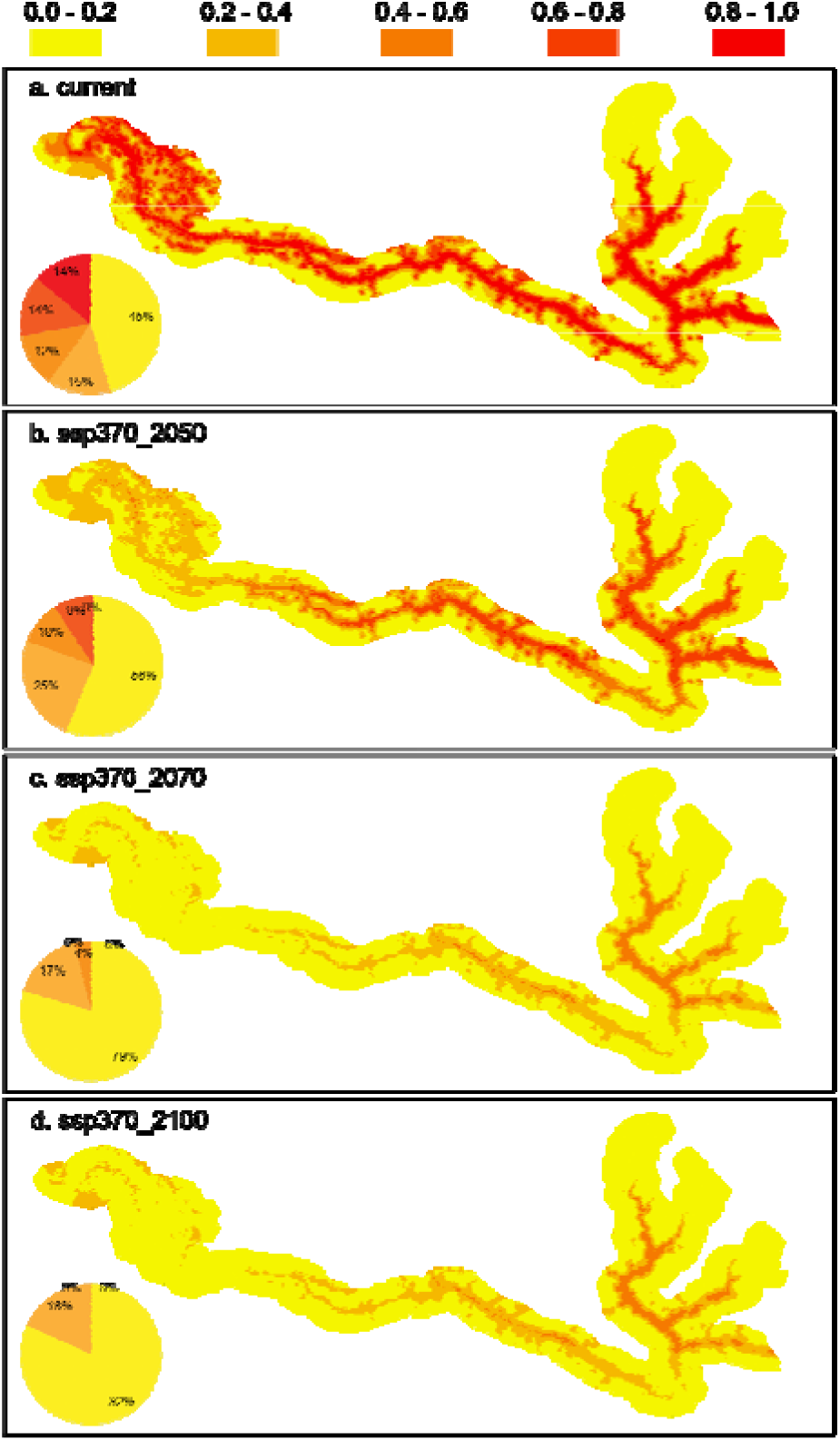
Spatial distribution of habitat suitability for the Sino-Mongolian beaver under current and future climate scenario SSP370. Maps illustrate the predicted habitat suitability for the Sino-Mongolian beaver: (a) under current climatic conditions, and projected suitability under the future Shared Socioeconomic Pathway (SSP) scenario SSP370 for (b) the 2050s, (c) the 2070s, and (d) the 2100s. The color gradient reflects the level of habitat suitability, where red indicates high suitability and yellow denotes low suitability.

Climatic habitat suitability decline was detected across all projected time periods under the SSP370 scenario (Fig. 7). From the current to the 2100s, 92% of the study area experienced a decline in suitability (Fig. 7a). Similarly, a large proportion of the area saw a decline from the present to the 2050s (84%, Fig. 7b). The situation of habitat suitability decline continued across subsequent periods: 95% of the area declined from the 2050s to the 2070s (Fig. 7c), and 70% declined from the 2070s to the 2100s (Fig. 7d). Similar patterns across SSP126 (Fig. S6) and SSP585 (Fig. S7) scenarios suggest the robustness of our analyses.

**Fig. 7.**
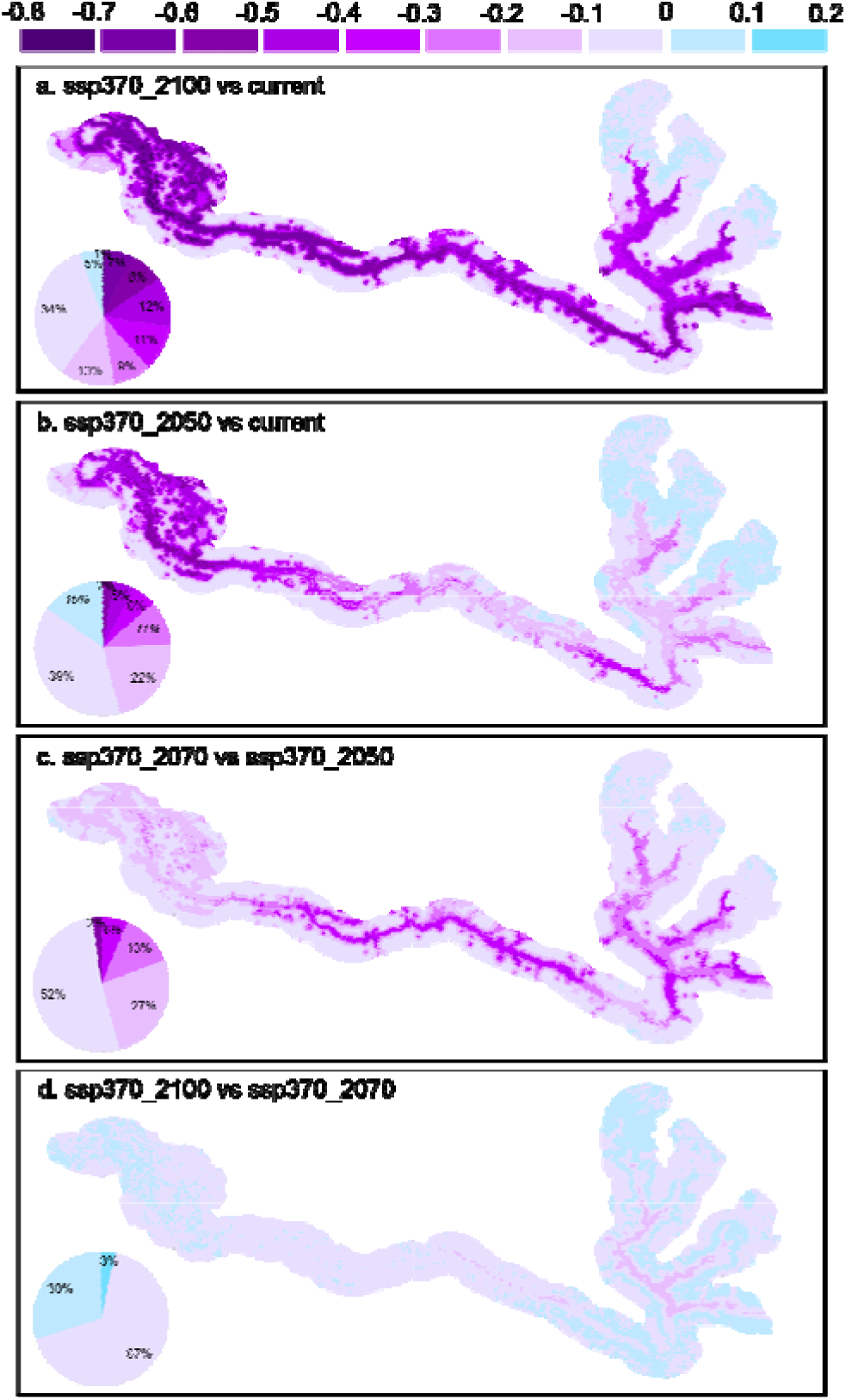
Temporal changes in habitat suitability for Sino-Mongolian beavers future climate scenario SSP370. Panels illustrate the projected difference in habitat suitability between specific time periods: (a) the 2100s relative to the current period; (b) the 2050s relative to the current period; (c) the 2070s relative to the 2050s; and (d) the 2100s relative to the 2070s. The color gradient indicates the magnitude of suitability change: purple represents an increase in suitability, while light blue represents a decrease.

## Discussion

### Population recovery: the success of phrased conservation model

Integrating our data in 14 systematic annual field surveys (2003-2023) with those in prior reports (1975-1989), this work reconstructed a long-term spatiotemporal population dynamic for the Sino-Mongolian beaver in China (Fig. 2). Our results confirmed a successful long-term beaver population recovery, a noteworthy conservation achievement largely attributing to the execution of a phased conservation strategy. Specifically, there was a close temporal coupling between the rate of beaver population recovery and the milestone conservation actions (Fig. 2). First, in late 1970s, administrations of Qinghe county started to forbid hunting activities on beaver. After that, the Xinjiang Burgen Beaver Nature Reserve was established in 1980. These initial conservation efforts promptly reversed the beaver population decline and promoted the population recovery from 27 colonies in 1975 to 93 colonies in 1983. Building on this initial success, the official release of the Key Protected Wildlife of China and the implementation of the Wildlife Protection Law in 1989 have further promoted the population recovery, rising up to 165 colonies by 1989 [16]. This early recovery trajectory, which was contingent upon nature reserve establishment and legislative protection, aligned well with the conservation success observed for the Eurasian beaver recovery in Europe [3, 6].

However, from 2003 to 2018, the beaver population entered a stable phase, characterized by temporal fluctuations (137-167 colonies) above a baseline established through earlier institutional efforts (Fig. 2). During this period, the stable yet fluctuating population recovery primarily resulted from enforced institutional efforts, such as the upgrade of the nature reserve in 2013. This enhancement not only stabilized the population size within the reserve but also fostered conservation efforts throughout the entire upstream area (Fig. 2). Subsequently, beaver population recovery accelerated markedly, rising from the pre-2018 plateau of 165 to 227 colonies by 2023. This accelerating trajectory highlights the critical role of multi-stakeholder collaboration, marked by a significant milestone in 2019, and encompassing contributions from governmental administration, conservation organizations, social support, and community-based conservation initiatives (SSCA). This paradigm shift, moving from exclusively restrictive protection toward inclusive co-management focused on securing social acceptance and mitigating human-wildlife conflicts [2, 37–40], successfully maximized conservation efficacy.

The efficacy of this phrased and collaborative conservation strategy in recent stage has been validated by the distinct spatial dynamics observed across different zones, specifically between the upstream and downstream segments and in areas within and outside the nature reserve (Fig. 2 and 3). Accompanying enhanced institutional conservation efforts, the upstream segment, which includes the nature reserve area, sustained relatively stable and high-density beaver populations (Fig. 3b, d and Fig. 4). By comparison, the downstream area, characterized by a higher anthropogenic disturbance pressure, experienced dramatic fluctuations. This downstream dynamic included a period of decline pre-2018 and a significant increase in colony counts and density post-2018 (Fig. 3-4 and S2). The population recovery in human-impacted landscapes in the downstream of Ulungur River parallels the contemporary Eurasian experience [41]. This successful outcome validates the application of the multi-stakeholder collaborative conservation model, which emphasize non-lethal mitigation and public outreach to foster human-beaver coexistence [42, 43], as pivotal for facilitating recovery in both population size and spatial expansion beyond core protected areas.

### The multi-dimensional complexity of habitat association

The relatively low accumulative variance explained by the first four components of PCA indicates the complexity of habitat association for the Sino-Mongolian beaver. The first component (PC1), accounting for only 23.7% of the total variance of nine local environmental variables (Fig. 5a), captured a critical topology and anthropogenetic disturbance gradient. Specifically, positive relationships with the distance to artificial dams, roads, and human settlements, coupled with negative relationships with elevation and NDVI, collectively suggest that beavers primarily colonize sites characterized by minimal human disturbance at low elevations (Fig. 5). This pronounced avoidance behavior represents a consistent ecological principle across the wider Eurasian range, where studies confirm that persistent anthropogenic pressures, such as infrastructure development, linear features, and habitat fragmentation [44–46], constitute the primary determinants limiting the range expansion of both Eurasian beaver and American beaver (*C. canadensis*). Furthermore, the negative association with distance to artificial dams highlights specific avoidance effect of beavers in regulated waterways [47]. Consequently, habitat association criteria appear heavily weighted toward human-beaver conflict avoidance [43, 45] rather than solely optimizing for resource availability. This indicates that conservation efforts must strategically manage or reduce the impact of surrounding direct and indirect human disturbance to ensure long-term viability and range expansion [48].

It is noteworthy that the high and negative effects of elevation and NDVI in PC1 partially reflect the complexity of habitat association and human-beaver conflicts in space. Specifically, the dense colonies in protected areas coincide with lower elevation (below 1000 m, Fig. S3). This suggests that elevation serves primarily as a proxy for the true ecological drivers, protection status and low anthropogenic pressure, rather than being a direct determinant of habitat suitability. Similarly, the observed correlation with the NDVI reflects habitat association at a landscape scale: most beaver nests lie in areas with relatively low NDVI, indicating that this pattern is likely a function of avoidance of highly vegetated human-dominated landscapes such as forest close to human settlements and roads, rather than a direct selection based on optimal forage or cover (Fig. S3f).

In contrast to macro-level anthropogenic constraints, the second component of the PCA (PC2, accounting for 13.2% of the total variance, Fig. 5a) reflect the essential hydro-geomorphic requirements and micro-habitat suitability, indicating that the species depends on specific physical and hydrological conditions [49]. For instance, the observed inverse correlations with water depth and nest height imply a selective preference for areas characterized by stable and optimal aquatic condition and adequate bank stability for secure, less exposed nest construction [50–52]. This preference for specific channel characteristics and flow stability underscores the importance of hydro-geomorphic integrity for beaver colonization [50–52]. These fine-scale resource associations [49] echo the established ecological requirements for *Castor* species across the Palearctic. The complexity of these inter-related factors confirms that effective habitat management requires a dual focus: minimizing anthropogenetic disturbance at the landscape scale while actively preserving the critical micro-habitat features that support beaver for successful colonization and reproduction.

We also believe that several factors are critical to beaver habitat selection, particularly biological stresses associated with livestock grazing, domestic dog disturbance, and biological invasions. Although livestock grazing and domestic dogs are prohibited within the nature reserve, a substantial proportion of beaver colonies occur outside the reserve, where they are exposed to competition for habitat and food resources as well as disturbance from livestock and domestic dogs [53]. In addition, recent studies suggest that two invasive semi-aquatic mammals, the American mink (*Neogale vison*) and the muskrat (*Ondatra zibethicus*), have established feral populations throughout the Ulungur River basin through escape from captivity or intentional release [54], which potentially increase habitat competition [55–57] and contribute to declines in beaver offspring survival [58, 59]. These factors were not explicitly incorporated into our analyses because they are closely associated with other indicators of human disturbance, such as distance to human settlements. Moreover, assessing their influence on beaver populations and habitat selection requires detailed data, such as continuous camera-trap records. We hope to address these aspects in future studies.

### Future vulnerability and adaptive management

Under future climate scenarios, our robust results of SDMs forecast a rapid and severe decline in habitat suitability for the Sino-Mongolian beaver across the study area. The limited high-suitability areas (about 14% of study areas) are projected to entirely or almost vanish by the 2050s, and consequently, the proportion of unsuitable habitat is expected to escalate to over 80% by the 2100s. This catastrophic decline is primarily driven by the high influence of climatic variables (Bio12, Bio3, Bio2, Bio4, and Bio19), indicating a fundamental vulnerability to changes in temperature and precipitation regimes [20]. As an arid-region obligate riparian species, the beaver’s reliance on stable water flow and depth makes it highly susceptible to climate-induced hydro-geomorphic instability [20]. Increasing aridity and extreme temperature variability directly diminish primary productivity, reduce food availability, and amplify the existing fragmentation of riparian corridors [60]. Consequently, the success achieved through current conservation efforts will be imminently threatened by emerging and pervasive climatic constraints under on-going climate change.

The projected vulnerability necessitates an urgent pivot toward climate-adaptive conservation, requiring the integration of SDM results into dynamic management planning. The foremost management priority must be water security and hydro-geomorphic stabilization. Effective management should focus on active water regulation to buffer against climate-induced flow variability, thereby ensuring stable water levels through targeted, non-lethal engineering measures [37, 45, 47, 48]. Secondly, efforts should concentrate on identifying and protecting climate refugia, specifically, the less-impacted riparian stretches that currently exhibit greater climatic resistance. These areas must be treated as critical source populations for future recolonization. Finally, to counteract the predicted habitat loss, intensive riparian restoration is required, focusing on enhancing the quality and connectivity of native vegetation along river segments identified by the models as retaining marginal suitability, thereby facilitating necessary dispersal and range shifts. This adaptive approach, combining large-scale climate buffering with landscape-scale human disturbance mitigating and fine-scale habitat enhancement, is essential to secure the long-term persistence of the Sino-Mongolian beaver in the face of escalating environmental change.

## Conclusion

In conclusion, this study demonstrates that the successful recovery of the Sino-Mongolian beaver is not monolithic but a phased outcome, first established by institutional efforts and subsequently accelerated by multi-stakeholder collaboration. Population dynamics demonstrate that the shift in conservation strategy, moving from purely restrictive protection toward social acceptance and conflict mitigation, was the critical driver enabling downstream range expansion. Ecological analyses reveal the complexity of habitat association: beavers primarily prioritize the avoidance of anthropogenic disturbance while simultaneously depending on specific hydro-geomorphic micro-environments for securing lodge and water stability. However, this hard-won conservation success faces an imminent and severe threat from climate change. SDMs project that climate shifts will lead to almost or complete disappearance of high-suitability habitats by the 2050s, posing a fundamental risk to this subspecies’ long-term persistence. Given these situations, future management strategies require urgent recalibration to adopt climate-adaptive conservation approaches. Key management recommendations include implementing engineering solutions to guarantee hydrological stability, identifying and strictly protecting high suitable areas as climate refugia to serve as critical source populations, and undertaking intensive riparian restoration in areas retaining marginal suitability to enhance habitat connectivity and enable beaver population expansion and reproduction.

## Acknowledgements

We especially thank Gang Chen from the Xinjiang Burgen Beaver National Nature Reserve Administration for his kind assistance with our field surveys. We also thank the following institutions and individuals for their support in this study: (1) Altay Prefecture Nature Conservation Association: Wenpu Huang, Dapeng Wang, Junliang Guo; (2) Altay Prefecture Forestry and Grassland Administration (formerly Altay Forestry Bureau): Rongguang Zheng, Yan Ge, Shan Lu, Weili Chang, Juncheng Tian, Feng Jiang, Bin Li, Xinchun Jiang, Mengk, Shuo Bai, Changliang Shao; (3) Xinjiang Burgen Beaver National Nature Reserve: Huaixi Feng, Yongshan Tao, Hai Rou, Jinlei Hou, Maerhabek, Chuang Huang, Peng Chen, Yuping Mao, Xuemin Jiao, Baoyu Wang, Daoken, Kekesogen, Ayimulat, Toha, Kerenbek, Qiakeer, Yerlan; (4) Qinghe County Forestry and Grassland Administration (formerly Qinghe County Forestry Bureau): Xinyong Xu, Renyuan Zhou, Wenqing Li, Zhumabek, Bahetibek Halier, Jiarheng, Tuoliubek, Danggele, Yesboli Ashmu, Bahetibek Bahdati, Wenzila, Hairula, Nuzhehmat, Halibek Rouhai, Bolati, Muerza, Wumuerbek, Aydar, Wunizila, Galin Bahar, Balafan Zhuodayi, Bolati Matai, Daohe Ganati, Nurboli Tuohatarkan, Telek Bulanbai, Helengbek Saimaihan, Nazharbek Huatbai, Hailati Hahada, Quhai, Saleck, Jinhu Li, Jiangsen Biduola, Yandong Qi, Xilong Du, Yerbole Jumahan, Xiangmin Liu, Wunizila Patihan, Hamitibek Galek, Jiekebai Tuoliwu, Haierola Aheti, Silamubek Heisha, Aydar Malik, Nurzhati Azatibek, Mahsuti Yerkengbek, Tuoliuwuhan Wunierbek, Alemjiang Awubabake, Pathan, Xingui, Nurahamaiti, Hanat, Baoxian Qiegeer, Mengxi Wu, Huanxibek Mawlanbai, Bazanbek Ers, Ganalbek Gafankeshi, Muzabek Hapura, Adari Jumadele, Humarbek Jiangsenhan, Asaharbek Humar, Tarafbek Galdebai, Mulatihan Aslam, Buyizhek Wuimudan, Jigerbek Sayasati, Ayiken Malhaba, Halibek Hapan; (5) Fuyun County Forestry and Grassland Administration (formerly Fuyun Forestry Bureau): Haibo Pu, Zhaoqing Yang, Bahexiabek, Tuleheng, Mulatihan, Dunbin Zhao, Talehati, Shanati, Huannixi, Huatibek, Manarbek, Yeurjiang, Tursunguli, Nurlanbek, Wumurzake, Hatila, Beshanbek, Aynur, Telewu; (6) Fuhai County Forestry and Grassland Administration (formerly Fuhai Forestry Bureau): Baohong Han, Xiaoyun Li, Wei Guo, Sailikebek, Tuerxun, Burulesibek, Adeli, Bahetibek, Kanghong Liu, Beken, Reahati, Hailati, Chunpeng Wei, Yandong Shan, Weiyanbek, Yerboli, Lazienbek, Nurlan, Adeliti, Hederbai, Ayihen, Jun Xie, Shalihen; (7) Buerjin County Forestry and Grassland Administration (formerly Buerjin Forestry Bureau): Jianabuer; (8) Xinjiang Altai Mountains Two-River-Source Nature Reserve: Tawu Mithati, Boken Manken; (9) Shandong University: Haoquan Lu, Jiayi Ji, Changqing Yu, Wen Shao, Manran Liu, Bujun Huang; (10) Xinjiang University: Dongzhi Liu, Congcong Du, Canxia Su, Pielizati, Yanqiu Chen, Xiangyu Duan, Wenming Wang. We also thank Man Qi, Hungwei Lin, and Wenyi Liu from Department of Biology, University of Oxford, for their constructive comments on early versions of manuscript.

This work was funded by National Key R&D Program of China (2024YFF1307500), National Natural Science Foundation of China (32270549), Altay Prefecture Nature Conservation Association (2019–2024), International S&T Cooperation Project of Xinjiang Uygur Autonomous Region (20136026), Central Government Capacity-Building Subsidy Projects (2014, 2015, 2016), Central Government Forest Ecological Benefit Compensation Fund, Central Government Project for the Rescue and Management of the Critically Endangered Sino-Mongolian Beaver (2021), the Third Comprehensive Scientific Expedition to Xinjiang (2021xjkk1200), National Forestry and Grassland Administration S&T Innovation Project (2022132017), and the Xinjiang Forestry Department S&T Project.

## Author contributions

The study was conceived and designed by Yuanbao Du, Kai Li, and Hongjun Chu; Wenwen Chu, Yingjie Qi, Chi Ma, Wenxu Lan, Xiaoyun Li, Rukeya Abulimit, Fangling Zheng, Ziyu Liu, Yuting Gao, Huanxin Liu, Chengbin He, Hongjun Chu actively participated in data collection; Yuanbao Du and Wenwen Chu conducted formal analysis and results visualization; Yuanbao Du and Wenwen Chu wrote the original draft, and all authors provided contributions to manuscript revision.

## Data Availability

The data that support the findings of this study are available from the corresponding authors upon reasonable request.

## Author statement

All authors declare that they have no conflicts of interest.

