## Supplementary material for "Looking backward for the future: long-term population recovery, habitat associations, and future climatic vulnerability of the critically endangered Sino-Mongolian beaver (*Castor fiber birulai*) in China": 02_Supple_Figures_Tables.docx

**Supplementary Materials**

**Supplementary Figures**

**Fig. S1.** **Colony size variation and key eco-engineering structures of the Sino-Mongolian beaver in northern Xinjiang, China.** Panels (a1)-(a5) illustrate the natural variation in beaver colony size, showing colonies containing 1 to 5 individuals. Key beaver-engineered structures are also shown: (b1) beaver nest; (b2) beaver-built dam; and (b3) food pile (winter food cache).

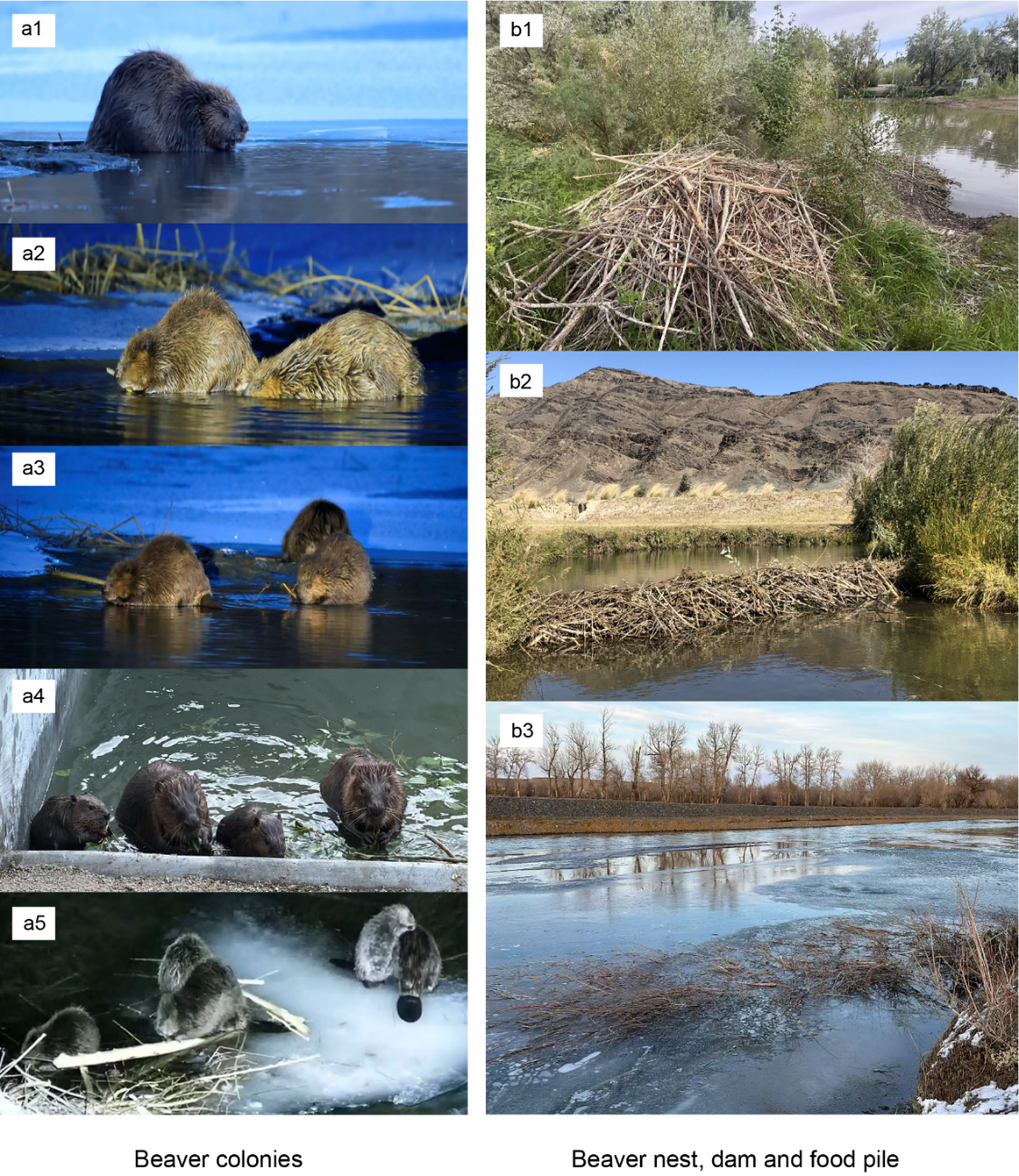

**Fig. S2.** **Spatial kernel density of beaver nest locations across five temporal stages.** Each panel displays the kernel density distribution of beaver nests, aggregated over a three-year period. The color gradient indicates nest density levels: dark purple-blue shows areas of high density, while light purple-blue represents areas of low density. Note that the beaver nest distribution data for 2021 was used in the density calculation for both stage 4 and stage 5.

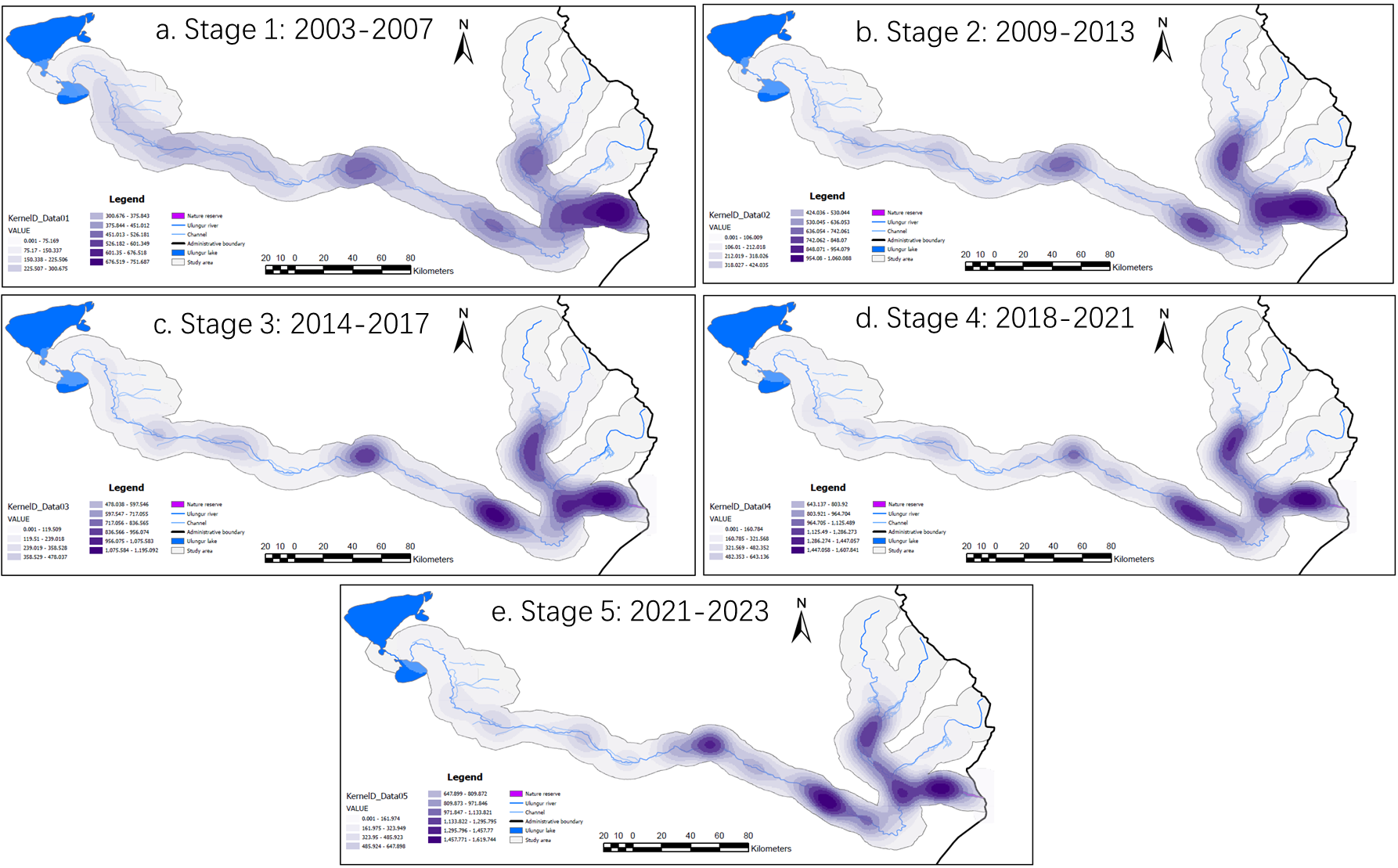

**Fig. S3. Density histograms of nine local-scale habitat variables for the Sino-Mongolian beaver.** The nine panels display the distribution (density histogram) of the following variables: (a) elevation; (b) distance to nearest road; (c) distance to nearest human settlement; (d) distance to nearest artificial dam; (e) distance to nearest river; (f) Normalized Difference Vegetation Index (NDVI) near the beaver nest; (g) water depth near the beaver nest; (h) forest canopy cover near the beaver nest; and (i) beaver nest height relative to the riverbank.

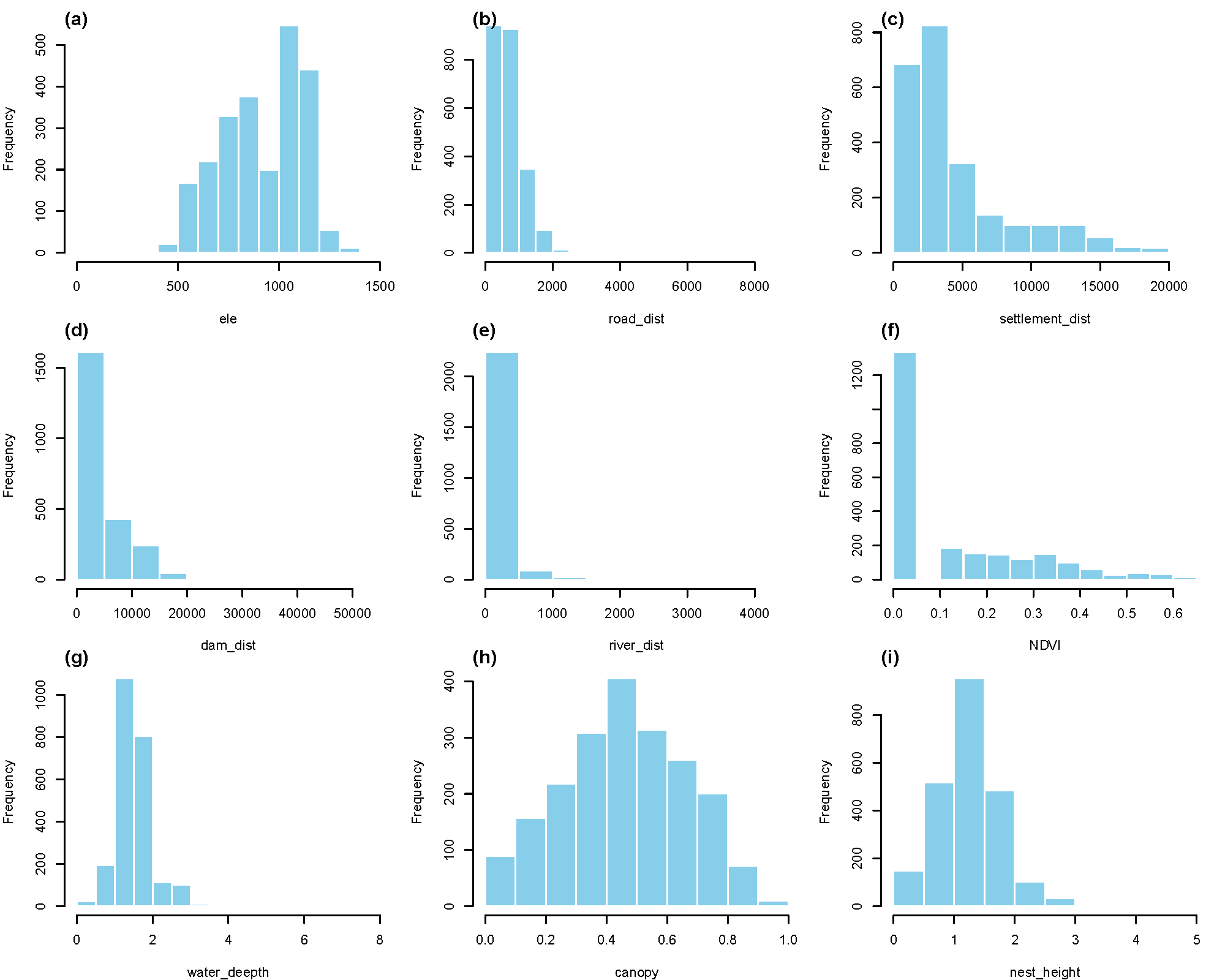

**Fig. S4.** **Spatial distribution of habitat suitability for the Sino-Mongolian beaver under current and future climate scenario SSP126.** Maps illustrate the predicted habitat suitability for the Sino-Mongolian beaver: (a) under current climatic conditions, and projected suitability under the future Shared Socioeconomic Pathway (SSP) scenario SSP126 for (b) the 2050s, (c) the 2070s, and (d) the 2100s. The color gradient reflects the level of habitat suitability, where red indicates high suitability and yellow denotes low suitability.

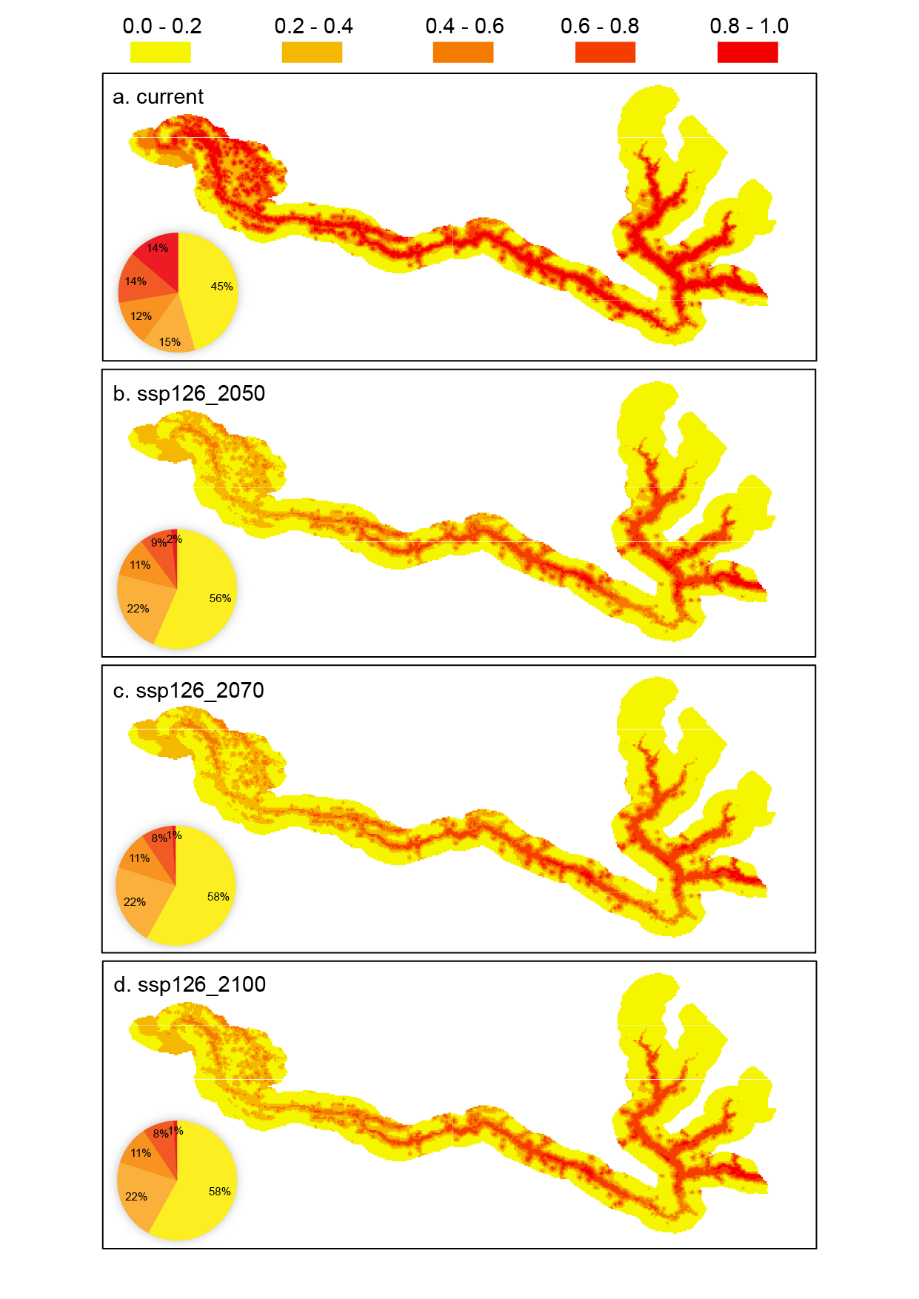

**Fig. S5.** **Spatial distribution of habitat suitability for the Sino-Mongolian beaver under current and future climate scenario SSP585.** Maps illustrate the predicted habitat suitability for the Sino-Mongolian beaver: (a) under current climatic conditions, and projected suitability under the future Shared Socioeconomic Pathway (SSP) scenario SSP585 for (b) the 2050s, (c) the 2070s, and (d) the 2100s. The color gradient reflects the level of habitat suitability, where red indicates high suitability and yellow denotes low suitability.

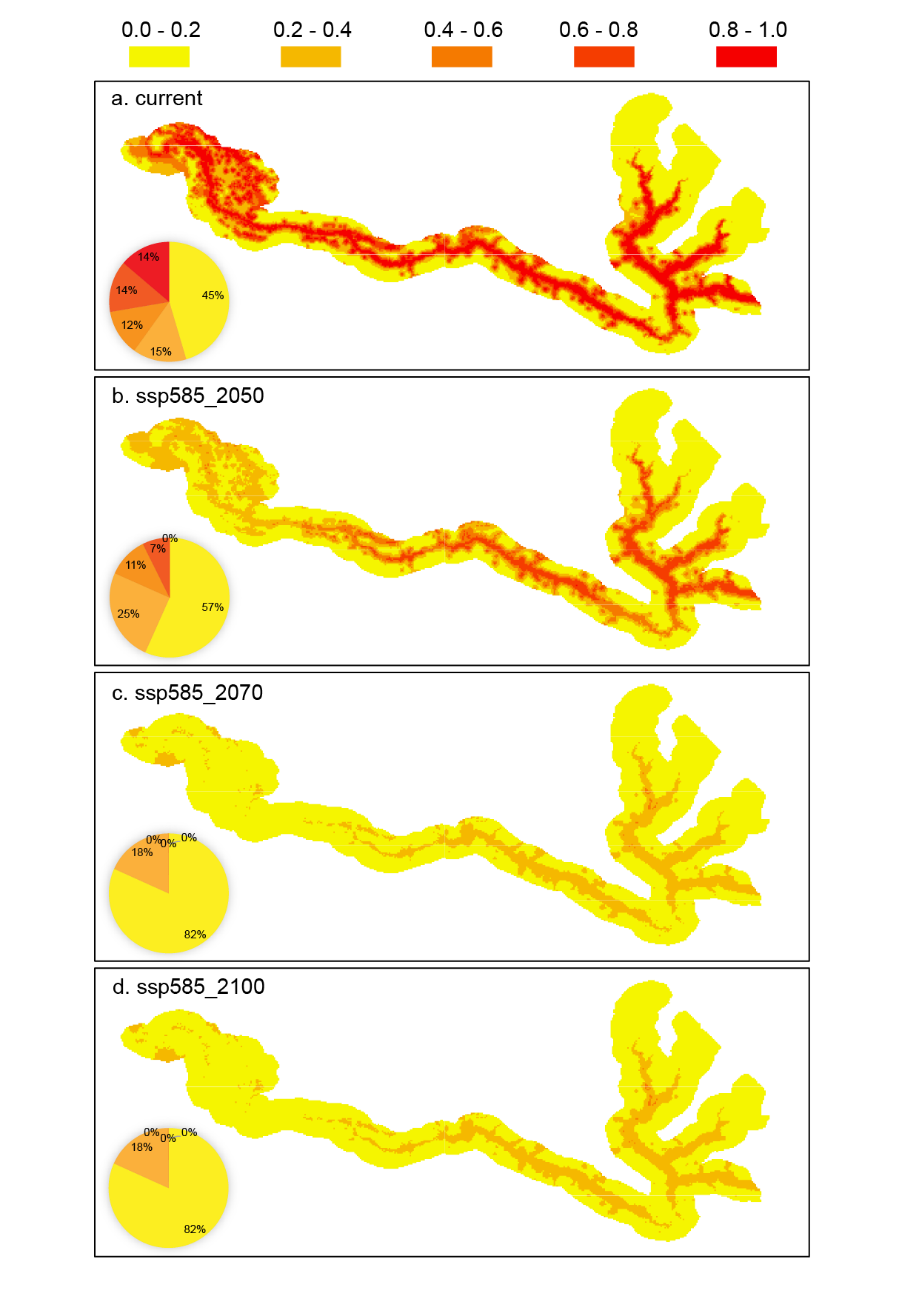

**Fig. S6.** **Temporal changes in habitat suitability for Sino-Mongolian beavers future climate scenario SSP126**. Panels illustrate the projected difference in habitat suitability between specific time periods: (a) the 2100s relative to the current period; (b) the 2050s relative to the current period; (c) the 2070s relative to the 2050s; and (d) the 2100s relative to the 2070s. The color gradient indicates the magnitude of suitability change: purple represents an increase in suitability, while light blue represents a decrease.

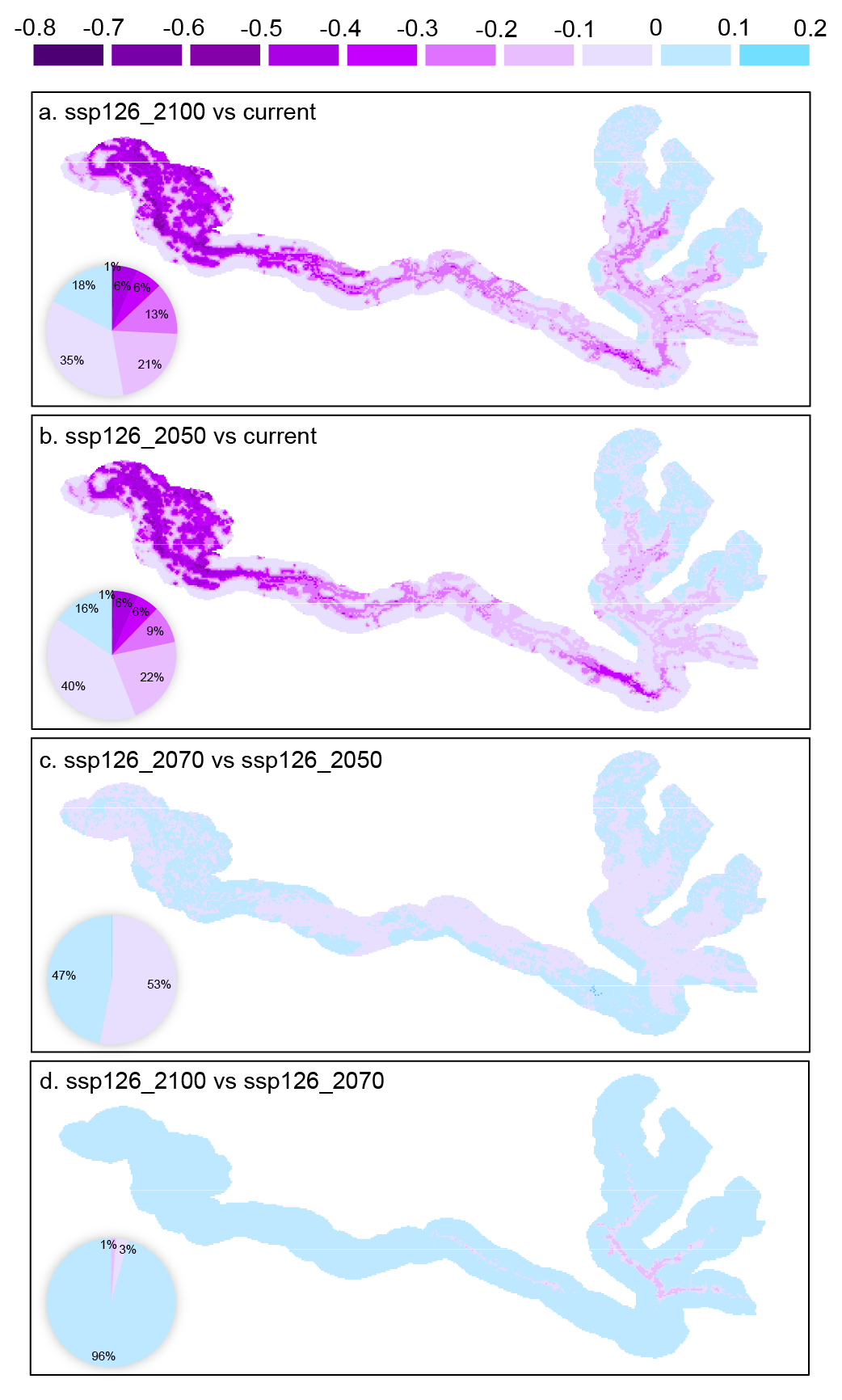

**Fig. S7.** **Temporal changes in habitat suitability for Sino-Mongolian beavers future climate scenario SSP585**. Panels illustrate the projected difference in habitat suitability between specific time periods: (a) the 2100s relative to the current period; (b) the 2050s relative to the current period; (c) the 2070s relative to the 2050s; and (d) the 2100s relative to the 2070s. The color gradient indicates the magnitude of suitability change: purple represents an increase in suitability, while light blue represents a decrease.

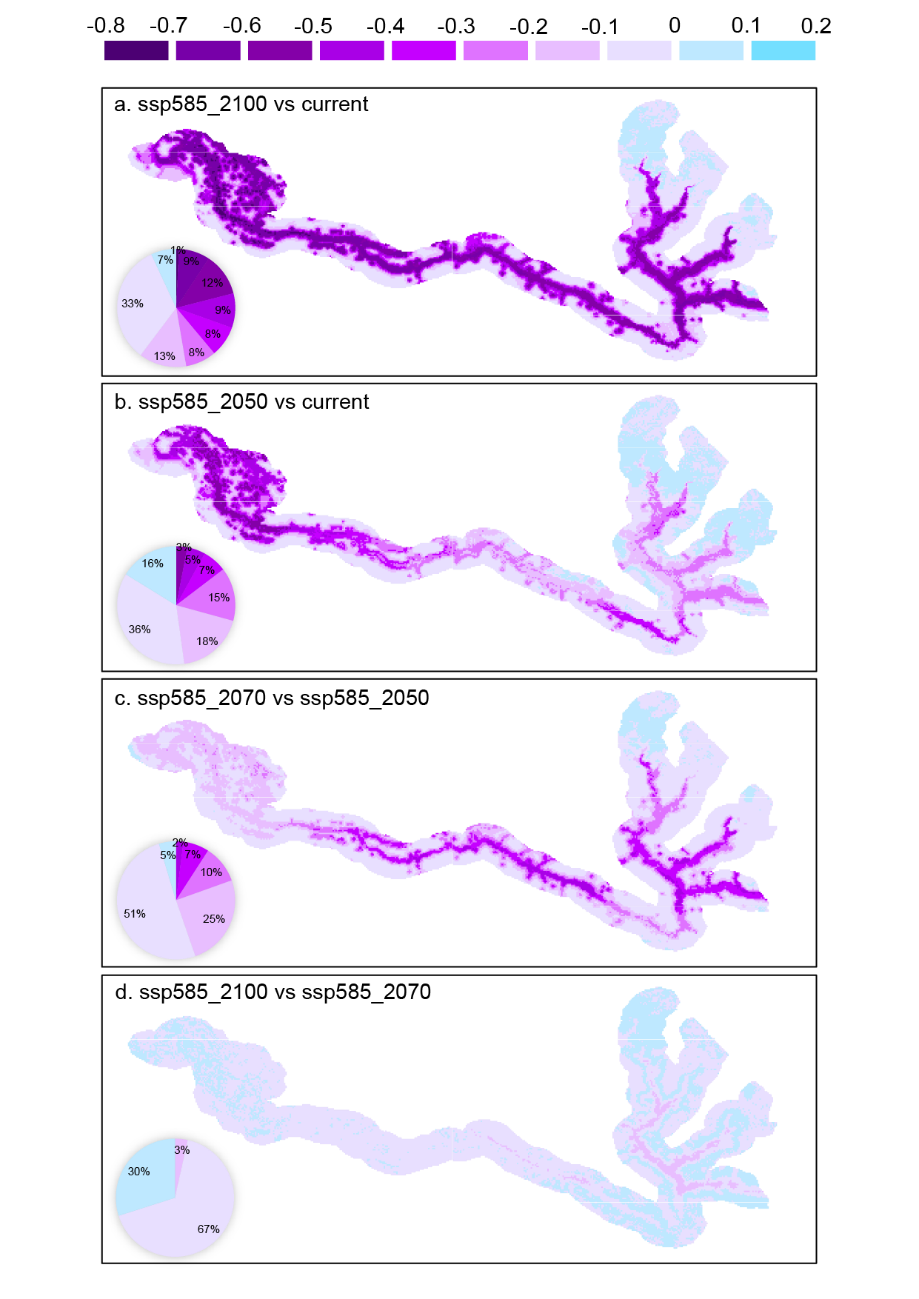

**Supplementary Tables**

**Table S1. Relative importance of principal components derived from the PCA analysis of nine habitat variables.**

|  | PC1 | PC2 | PC3 | PC4 | PC5 | PC6 | PC7 | PC8 | PC9 |
| --- | --- | --- | --- | --- | --- | --- | --- | --- | --- |
| Standard deviation | 1.461 | 1.090 | 1.033 | 1.012 | 0.975 | 0.947 | 0.889 | 0.783 | 0.573 |
| Proportion of Variance | 0.237 | 0.132 | 0.118 | 0.113 | 0.105 | 0.099 | 0.087 | 0.068 | 0.036 |
| Cumulative Proportion | 0.237 | 0.369 | 0.487 | 0.601 | 0.707 | 0.807 | 0.895 | 0.963 | 1 |

**Table S2. Species distribution model validation metrics: AUC, TSS, and corresponding thresholds.**

| Model_ID | AUC | TSS | threshold |
| --- | --- | --- | --- |
| 1 | 1 | 0.995 | 0.53365 |
| 2 | 1 | 0.998 | 0.598883 |
| 3 | 1 | 0.996 | 0.531783 |
| 4 | 1 | 0.996 | 0.67895 |
| 5 | 0.999 | 0.99 | 0.670333 |
| 6 | 1 | 0.992 | 0.5006 |
| 7 | 1 | 0.998 | 0.616933 |
| 8 | 1 | 0.995 | 0.6426 |
| 9 | 1 | 0.992 | 0.695767 |
| 10 | 0.999 | 0.993 | 0.468933 |
| 11 | 0.993 | 0.942 | 0.480468 |
| 12 | 0.991 | 0.959 | 0.410462 |
| 13 | 0.991 | 0.945 | 0.767964 |
| 14 | 0.995 | 0.964 | 0.514119 |
| 15 | 0.995 | 0.957 | 0.716698 |
| 16 | 0.99 | 0.933 | 0.671858 |
| 17 | 0.991 | 0.954 | 0.722678 |
| 18 | 0.992 | 0.963 | 0.713574 |
| 19 | 0.996 | 0.968 | 0.594578 |
| 20 | 0.989 | 0.951 | 0.676272 |
| 21 | 0.999 | 0.981 | 0.498337 |
| 22 | 1 | 0.992 | 0.449339 |
| 23 | 0.999 | 0.981 | 0.484785 |
| 24 | 0.998 | 0.991 | 0.431226 |
| 25 | 0.998 | 0.987 | 0.433716 |
| 26 | 0.999 | 0.982 | 0.485958 |
| 27 | 0.998 | 0.989 | 0.446454 |
| 28 | 0.999 | 0.989 | 0.316473 |
| 29 | 0.996 | 0.983 | 0.366741 |
| 30 | 0.997 | 0.985 | 0.446262 |
| Mean±SD | 0.997±0.004 | 0.978±0.019 | 0.552±0.121 |

**Table S3. Relative importance of predictive variables in the SDMs.** The table shows the relative importance of nine predictive variables in the SDM. 'Cor_test' represents the mean correlation coefficient between the variable's importance and the model's performance, with 'Cor_lower' and 'Cor_upper' denoting the lower and upper confidence boundaries, respectively. 'AUCtest' represents the AUC-based relative importance for each variable, with 'AUCtest.lower' and 'AUCtest.upper' indicating the corresponding confidence boundaries. Bio2–Bio19 represent six bioclimatic variables. settlement_dist is the nearest distance to human settlements (derived from the Global Human Settlement Layer). river_dist is the nearest distance to rivers, lakes, and channels (derived from the Global Lakes and Wetlands Database). road_dist is the nearest distance to roads (derived from the Global Roads Database).

| Predictors | Cor_test | Cor_lower | Cor_upper | AUCtest | AUCtest.lower | AUCtest.upper |
| --- | --- | --- | --- | --- | --- | --- |
| Bio2 | 0.178 | 0.010 | 0.345 | 0.120 | -0.004 | 0.244 |
| Bio3 | 0.253 | 0.041 | 0.466 | 0.203 | 0.004 | 0.402 |
| Bio4 | 0.162 | 0.061 | 0.264 | 0.095 | 0.005 | 0.185 |
| Bio6 | 0.084 | 0.057 | 0.111 | 0.031 | 0.008 | 0.054 |
| Bio12 | 0.274 | 0.179 | 0.369 | 0.203 | 0.062 | 0.343 |
| Bio19 | 0.173 | 0.134 | 0.211 | 0.091 | 0.047 | 0.136 |
| settlement_dist | 0.058 | 0.023 | 0.094 | 0.018 | 0.002 | 0.033 |
| river_dist | 0.159 | 0.116 | 0.202 | 0.054 | 0.023 | 0.085 |
| road_dist | 0.011 | 0.002 | 0.019 | 0.002 | 0.000 | 0.003 |
